# Privacy-Preserving and Robust Watermarking on Sequential Genome Data using Belief Propagation and Local Differential Privacy

**DOI:** 10.1101/2020.09.04.283135

**Authors:** Abdullah Çağlar Öksüz, Erman Ayday, Uğur Güdükbay

## Abstract

**Motivation:** Genome data is a subject of study for both biology and computer science since the start of Human Genome Project in 1990. Since then, genome sequencing for medical and social purposes becomes more and more available and affordable. Genome data can be shared on public websites or with service providers. However, this sharing compromises the privacy of donors even under partial sharing conditions. We mainly focus on the liability aspect ensued by unauthorized sharing of these genome data. One of the techniques to address the liability issues in data sharing is watermarking mechanism.

**Results:** To detect malicious correspondents and service providers (SPs) -whose aim is to share genome data without individuals’ consent and undetected-, we propose a novel watermarking method on sequential genome data using belief propagation algorithm. In our method, we have two criteria to satisfy. (i) Embedding robust watermarks so that the malicious adversaries can not temper the watermark by modification and are identified with high probability (ii) Achieving *ϵ*-local differential privacy in all data sharings with SPs. For the preservation of system robustness against single SP and collusion attacks, we consider publicly available genomic information like Minor Allele Frequency, Linkage Disequilibrium, Phenotype Information and Familial Information. Our proposed scheme achieves 100% detection rate against the single SP attacks with only 3% watermark length. For the worst case scenario of collusion attacks (50% of SPs are malicious), 80% detection is achieved with 5% watermark length and 90% detection is achieved with 10% watermark length. For all cases, *ϵ*’s impact on precision remained negligible and high privacy is ensured.

**Availability:** https://github.com/acoksuz/PPRW_SGD_BPLDP

## 1 Introduction

Digital watermarking is one of the most important technological milestones in digital data hiding. It is used as a technique to hide a message or pattern within the data itself for various reasons like copyright protection or source tracking of digitally shared data. Watermarks may contain information about the legal owner of data, distribution versions, and access rights (Lee and Jung, 2001). Although watermarking has a wide range of applications, implementation schemes require different configurations for each use case and data type. For embedding copyright information in data and source tracking, robustness (Barni and Bartolini, 2004) against modifications is the crucial factor to preserve. The factors influencing such configurations alter depending on the characteristics of data, as well. Nonexistent redundancy, the existence of correlations and prior knowledge for inference, and the utility’s impact on attacker inference in sequential data are such factors that prevent the explicit implementation of digital data watermarking methods on sequential data.

We propose a novel watermarking scheme for sharing sequential genomic data consisting of Single Nucleotide Polymorphism (SNPs) with three states to be used by medical service providers (SPs). Each SP has access to the uniquely watermarked version of some individuals’ genomic data. The preliminaries expected from this watermarking scheme are robustness against watermark tampering attacks such as modification and removal, imperceptibility for not revealing watermark locations, preserving utility of original data to a degree through a minimum number of changes and satisfy local differential privacy in watermarks so that the watermarked versions of the data are indistinguishable from actual human genomic data and provide plausible deniability to data owners. By doing so, the watermarked data will not be shared in an unauthorized way and the source(s) of a leak will be easily identified by the data owner. To solve this multi-objective optimization problem, we use the belief propagation algorithm, which helps us to determine optimal watermarking indices on data that may preserve robustness with the highest probabilities. In belief propagation we consider the public knowledge about the human genome like Minor Allele Frequencies (MAFs) of SNPs, point-wise correlations between SNPs, called Linkage Disequilibrium (LD) and the prior knowledge of genotype and phenotype information that may potentially leak probabilities about watermarked points. Through conversion of prior information (MAF, LD, and so on) into the marginal probability distribution of the three SNP states, we manage to infer the state probabilities of each SNP. Our contributions are as follows:

1. We introduce a novel method for watermarking sequential data concerning the privacy of data and the robustness of watermark at the same time. We present the method’s strengths and weaknesses in various attack scenarios and provide insight into the weaknesses.
2. Our method uses prior information (MAFs, phenotype information, and so on) and inherent correlations to infer the state probabilities of SNPs. Using these inferred probabilities, we select SNPs that satisfy the following two criteria in a non-deterministic setup: a low probability of robustness decrease (change resistance) when attacked and a low utility loss (efficient index selection) when changed. By giving priority to these SNP points for watermarking, we guarantee the preservation of robustness and utility in data against various attacks. Besides, the identification probabilities of single SNPs using prior information are decreased with this method.
3. We test the robustness and limitations of our method using collusion (i.e., a comparison using multiple watermarked copies of the same data) and modification attacks and demonstrate how to reach a high probability of detection with various parameters, such as the watermark length, the number of SPs, the number of malicious SPs and the *ϵ* coefficient of local differential privacy.
4. We introduce randomly distributed non-genome-conflicting noise generated for the data to act naturally as watermarks and create imperceptible watermark patterns from the normal human genome if not attacked with collusion. Hence, rather than creating a fixed number of point-wise changes and tracking these changes for source tracking, we evaluate the whole data and reach a high probability of detection with a minimum number of changes.
5. We introduce watermarking schemes that satisfy *ϵ*-local differential privacy and plausible deniability in data along with it for data owners who value additional manners of enhanced privacy.

We provide a summary of related background on genomics, specifically Minor Allele Frequency (MAF) and Linkage Disequilibrium (LD), and Local Differential Privacy (LDP), in Appendix, § 1.

## 2 Related Works

Recent advances in molecular biology and genetics and next-generation sequencing increased the amount of genomics data significantly (Carter, 2019). While achieving a breakthrough in the genomics field, genomics data posses an important privacy risk for individuals by carrying sensitive information, i.e., kinship, disease, disorder, or pathological conditions (Grishin *et al*., 2019). Thus, collecting, sharing, and conducting research on the genomic data became difficult due to privacy regulations (CMS, 1996). Further, Humbert *et al*. (2013) show that sharing genomic data also threatens the relatives due to kin relation of genomic data. To this end, several works have been conducted to find emerging ways of privacy-preserving collection and analysis of the genomic and medical data in the last decade. Some of the privacy-preserving techniques used for medical data collection are k-anonymity, l-diversity, de-identification, perturbation, anonymization, or t-closeness (Samurai and Sweeney, 1998; Machanavajjhala *et al*., 2007; Wylie and Mineau, 2003; Kargupta *et al*., 2003; Li and Li, 2006). These methods, however, provide limited privacy protection and are prone to inference attacks. Ayday *et al*. (2013) proposed obfuscation methods in which the output domain is divided into several sections and one section is reserved for genomic data protection.

Digital watermarking is a technique usually used for copy protection by inserting a pattern to the digital signal such as song, image, or video (Cox *et al*., 2008). It is an attack counter-measure for the case of leakage or sharing without consent. Watermarking does not prevent leakage; it is used as a detection technique for malicious parties. Some of the application areas of digital watermarking are images (Huo and Gao, 2006), audio signals (Asad *et al*., 2011) and text documents (Topkara *et al*., 2006). These methods are prone to collusion attacks where malicious parties collude with other parties to detect the watermark.

Watermarking schemes for sequential data, especially for genomic data, are very rare. Kozat *et al*. (2008) proposed a steganography-based watermarking scheme for sequential electrocardiography data to hide private meta-data. Iftikhar *et al*. (2015) proposed a robust and distortion-free watermarking scheme, *GenInfoGuard*, for genomic data. They use features selected from the data for watermark embedding. Similar to ours, Liss *et al*. (2012) proposed a permanent watermarking scheme in synthetic genes that embeds binary string messages on open-frame synonymous amino-acid codon regions. Heider and Barnekow (2008) proposed the use of artificial dummy strands to act like watermarks on DNA.

Ayday *et al*. (2019) proposed a robust watermarking scheme for sharing sequential data against potential collusion attacks by using non-linear optimization. Our objective model is similar to theirs. However, different from their study, we consider the additive type of prior information scheme in which besides correlations, all sequential genomic data related information like familial genomes, phenotype states can be included by using factor nodes in belief propagation algorithm. Besides, we designed single SP and collusion attacks that incorporate all the information that can be gained from the correlation attacks and from one another within so that the worst-case scenarios are assumed and the attack model becomes more inclusive. Another difference between our method and theirs is the incorporation of *ϵ*-local differential privacy as an extra measure of privacy without impacting security. Andrés *et al*. (2012) proposed a method of embedding noise in sequential location data for geo-indistinguishability without violating the differential privacy. Inspired from their study and their new differential privacy criteria, we implemented a local set up so that extra criteria in the watermarking process checks every data index against the differential privacy violations locally and prevents the violating versions from getting shared.

## 3 Problem Definition

We present the data, system, and threat models, and the objective of our system. Frequently used symbols and notations are presented in Table 1.

**Table 1.** Frequently used symbols and notations

|  |  |
| --- | --- |
| $x_1, \dots, x_{d_l}$ | Set of data points |
| $y_1, \dots, y_m$ | Possible values (states) of a data point |
| $d_l$ | Length of the data |
| $w_l$ | Length of the watermark |
| $h$ | Total number of SPs |
| $I_k$ | Index set of data points that are shared with $k^{th}$ SP |
| $J_k$ | Index set of data points that are watermarked for $k^{th}$ SP |
| $Z_k$ | Watermark pattern of $k^{th}$ SP |
| $W_k$ | Watermarked data shared with $k^{th}$ SP |
| $\epsilon$ | Local differential privacy coefficient |
| $S_{I_i}^k$ | Set of states for index $i$ that are shared with the first $k$ SPs |

### 3.1 Data Model

Sequential data contain ordered data points *x*_1_, *x*_2_, …, *x_d_l__*, where *d_l_* is the length of the data. The values of *x_i_* can be in different states from the set {*y*_1_, *y*_2_, …, *y_m_*} depending on the type of the data. For example, *x_i_* can be an hour, minute, or second triplets ranging from 0 to 23, 59, 59, respectively, for timestamp data. For our system, we will use 0, 1, and 2 for the SNP states of *homozygous major, heterozygous*, and *homozygous minor*, respectively. The length of the data is *d_l_* and the number of points that will be watermarked at the end of the algorithm is *w_l_*.

### 3.2 System Model

We consider a system between data owner (Alice) and multiple service providers (SPs) with whom Alice shares sequential data, e.g., human genome, text, and location data. For text, the SP can be any service provider working on Natural Language Processing. For genome, service providers can be medical researchers, medical institutions, or bio-technical companies. Alice may decide to share the whole data or parts of it to receive different services. Besides, the parts shared with each SP may differ.

For all cases listed above, Alice wants to ensure that her data will not be shared unauthorized by SPs. If the data is shared anyway, she wants to preserve a degree of differential privacy and detect the malicious SP(s) who shared the data. She uses watermarking and shares a watermarked version of the data, which satisfies the degree of privacy she desires. These versions are produced through removing of certain parts or modifying the data. Data indices most optimal for satisfying the criteria given above should be calculated beforehand by considering the structure, distribution, and vulnerabilities of the data. To calculate the complex probability distributions of multi-variable sequential data that satisfy Alice’s demands, we use Belief Propagation (BP). Please confirm Appendix § 2 for the details of BP and our proposed solution. Other graph inference methods could have been used for the calculations but BP is adapted because of its fast approximation efficiency in non-loopy graph networks.

Watermarking is mostly done by changing the status of data indices. Adding dummy variables is an example of methods that do not change the actual values but common methods used for watermarking are usually removal or modification. Since a slight addition in sequential data causes a shift in other indices, it impacts the rest of the retrieval and embedding processes like the butterfly effect. We stick with the watermarking method by removal or modification. In a broader sense, non-sharing can be considered as modifying the status of a certain index into “non-available.” Normally, the security of a watermarking scheme increases along with the length of the watermark against attacks. However, a robust watermark should be short and as efficient as possible to maximize the detection probability of malicious SPs without reducing utility significantly.

Malicious SPs try to lower their chance of getting detected while leaking the data. If the system cannot identify the source of leakage due to various attacks, SPs avoid getting caught. To do so, SPs tamper the watermarks via the same processes of embedding: by removal or modifications. Sharing a portion of data is an example of removal to avoid detection but this reduces the amount of information data contains. On the other hand, if the watermarked indices are known or inferred, changing the values of watermarked states rather than albeit removing them will help SPs to share the data undetected. Watermarked indices on the data can be found by the collaboration of multiple SPs who compare their versions of the data with each other by a collusion attack. Another method for finding the indices is using prior knowledge to infer the actual states of the data and looking for discrepancies by a single SP attack. Therefore, the belief propagation algorithm helps us to find the optimum indices that are vulnerable to these attacks very little and satisfy the conditions of maximized probability of detection against various attacks, ensured privacy and minimum utility loss.

### 3.3 Threat Model

The objective of our proposed system and watermarking, in general, is identifying the source(s) of leakage when the data is shared unauthorized. In the threat model, contrary to our objective, the goal of malicious SPs is to share the data undetected. This goal can be achieved by decreasing the robustness of the watermark which prevents the identification of the leakage source(s). Malicious SPs can identify high probability watermark points and tamper the watermark pattern by removal or modification. For such scenarios, we presume that malicious SPs will not do blind attacks without the prior information of watermarked indices. These types of attacks will decrease the utility of data more than the robustness of the watermarking scheme and render the data useless. Hence, we introduce two attack models that incorporates the additional one within both based on probabilistic identification that test the robustness of the watermarks that our proposed method generates.

#### Single SP Attack

In this attack, a single malicious SP is expected to use the prior information available to infer the actual states of the data and identify the watermarked indices without collaborating with other SPs. Examples of prior information include *MAFs*, *genotype* and *phenotype* information of parents. For each data point, malicious SP finds the posterior probability of each state given the prior information *Pr*(*x_i_* = *y*| *prior information*) and compares it with the expected probability of given state *x_i_* = *y, y* ∈ {*y*_1_, *y*_2_, *y*_3_, …, *y_m_*}. If the difference between the posterior and expected probabilities for the given state is high, it may be an indication of a watermarked index. We assume that the malicious SP knows the watermark length *w_l_*, hence SPs select the top *w_l_* indices with the highest differences in probability as watermarked and implement an attack.

Another vulnerability that malicious SPs can exploit is the inherent correlations and their values in the data to infer the actual states of correlated indices. For genomic data, *linkage disequilibrium* (*LD*); nonrandom association of certain alleles is an example of such correlations. LD is a property of certain alleles; not their loci. The correlation of alleles {*A*, *B*} in loci {*I_A_*, *I_B_*} will not hold if either *A* or *B* changes. The asymmetric correlation observed in LD is a valid method of representation for other sequential data types. We treat the correlations in the data as pairwise and asymmetric in the proposed system.

#### Collusion Attack

In addition to the knowledge obtained via a single SP attack, multiple SPs that receive the same proportion of data can vertically align their data to identify watermarked points. When SPs align their data, there will be indices with different states that can be considered as definitely watermarked. The proportion of data shared with SPs may differ, which will decrease the efficiency of alignment. However, for the construction of a strong model against worst-case scenarios, the system considers the same data is shared among all SPs. Potentially watermarked indices received from collusion attack can be used along with prior information obtained from running a single SP attack and this attack type detects further more watermarked indices than the single SP attack.

## 4 Proposed Solution

When Alice wants to share her data with *SP_i_*, they employ the following protocol. The *SP_i_* sends a request to Alice providing the indices required from her data, denoted as *I_i_*. Then, Alice generates a list of available indices most suitable for watermarking *J_i_* that satisfies *J_i_* ⊂ *I_i_* and | *J_i_* | = *w_l_*. *J_i_* is generated by the BP-based watermarking algorithm, which will be discussed in detail in the sequel. Finally, Alice inserts watermark into the indices of *J_i_*. If the data is in binary form, it is as simple as changing 0 to 1 or vice versa. Otherwise, for the given state *x_i_*, a different state *y_i_* from the set *y_i_* ∈ {*y*_1_, *y*_2_, …, *y_m_*} and *y_i_* ≠ *x_i_* is chosen to be a part of the watermark pattern. In non-binary selection, if the given index contains correlation with other indices, the selection is determined by the probabilities and statistics of the correlated indices so that the watermark would not be vulnerable to the correlation attacks. Otherwise, it is a random selection with uniform distribution.

Our method relies on the BP algorithm that uses prior information and previous shared versions of the data to identify the indices with maximized detection probabilities of malicious SPs, ensured privacy and minimized utility loss when modified for watermarking. BP is an iterative message-passing algorithm used for the inference of unobserved random variables from the observed ones. We use this algorithm to infer the probability distributions of indices given the multi-variable prior information, attack scenarios, and privacy criteria. Normally, the factorization of prior information marginal probabilities could be used for a part of the inference of state probabilities. However, probability calculation gets exponentially complex as the dimensions of the data and the variety of prior information increases. Because BP approximates the actual state probabilities in a finite number of iterations, it is much more efficient than factorized calculation. The main idea is to represent the probability distribution of variable nodes by factorization into products of local functions in factor nodes.

### 4.1 Nodes and Messages

We describe the general setup and the details of the proposed BP-based algorithm for genomic data. BP consists of factor nodes, variable nodes, and messages between them. Connections between variable nodes and factor nodes are given in the factor graph (see Figure 1).

**Fig. 1.**
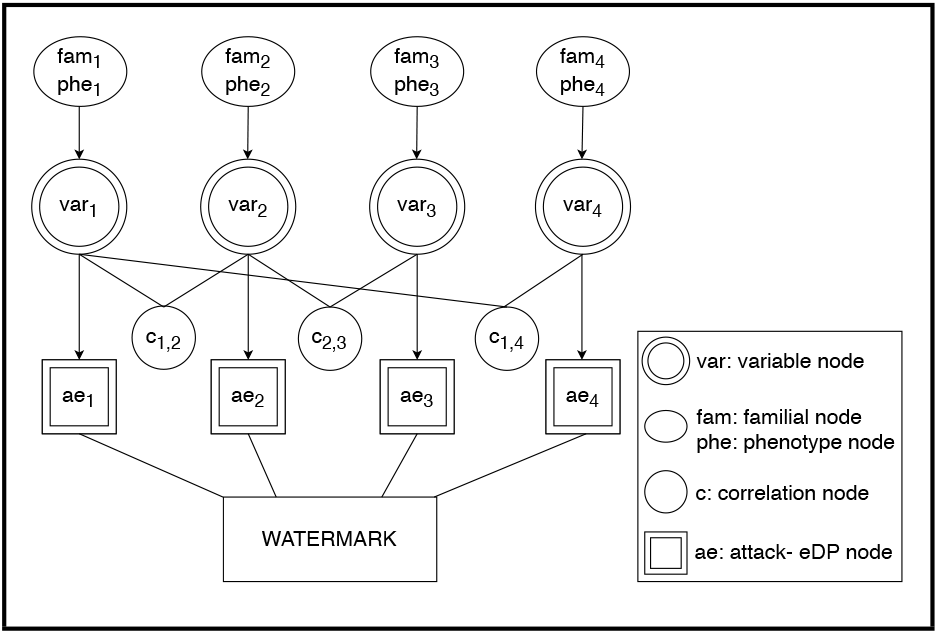
Factor graph representation of variable nodes and attack-eLDP interactions with other factor nodes: familial nodes, phenotype nodes, and correlation nodes.

The notations for the messages at the *v^th^* iteration are as follows:

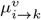: Message from variable node *var_i_* to factor or attack and *ϵ*-local differential privacy (*attack-eLDP*) node *k*.
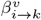: Message from familial node *fam_i_* to variable node *k*.
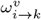: Message from phenotype node *phe_i_* to variable node *k*.
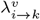: Message from correlation node *c_i,k_* to variable node *k*.
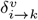: Message from *attack-eLDP* node *ae_i_* to be used as parameters for the watermarking algorithm.

#### 4.1.1 Variable Nodes

Variable (decision) nodes represent unknown variables. Each variable node sends and receives messages from factor nodes to learn and update its beliefs. Its purpose is to infer the marginal state probabilities of all indices that can be obtained from prior information. For genomic data, this information is the publicly known statistics, such as linkage disequilibrium (LD) correlations, familial genomic traits, and phenotype features.

For each node, we have a marginal probability distribution of states *y*_1_, *y*_2_, …, *y_m_*. Each variable node, *var_i_*, represents the marginal probability distributions of the *i^th^* unknown variable in the format of [*P*(*x_i_* = *y*_1_), *P*(*x_i_* = *y*_2_), …, *P*(*x_i_* = *y_m_*)] so that each *P* corresponds to the probability of one y and all sums up to 1. The probability distributions in variable nodes are calculated by multiplying the probability distributions coming from the neighboring factor nodes, such as correlation, familial, and phenotype nodes. The message 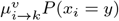 from variable node *i* to factor node *k* indicates that *P*(*x_i_* = *y*) at *v^th^* iteration where *y* ∈ {0, 1, 2}. Equation 1 provides the function for the representation of a message from variable node *i* to correlation factor node *k*:

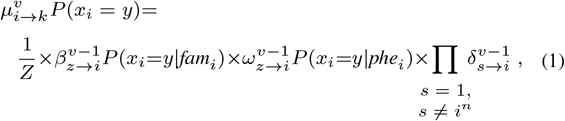

where *Z* is a normalization constant and 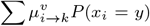 for all *y* must be equal to 1.

#### 4.1.2 Factor Nodes

Factor nodes represent the functions of factorized joint probability distributions of variable nodes. The messages received or sent by them might be dependent on multiple variable nodes as well as a single variable node. Factor nodes might also be independent and fixed from the start. For genomic data, the correlation between SNPs (LD) can be given as the example of the first case. In such scenario, variable node *var_i_* is connected to a correlation factor node *c_i,j_* along with the correlated variable node *var_j_*. For the second case of dependency on a single variable, a message passed into the AE-node is determined by the current state of any variable node can be given as an example. For the third case of dependency, family genomic information predetermined from the start can be given as an example. Let us assume for an SNP *x*, genomic information obtained from the family (father and mother) of certain individual *L* is *x_L,f_* = 0 (homozygous major) and *x_L,m_* = 1 (heterozygous). Then, we can safely predict the marginal probability distribution of that individual’s SNP as *P*(*L*, *x*)=[0.5 0.5 0] using the Mendelian Law of Segregation. This probability distribution is constant and not dependent on any value that the variable node might get. Therefore, throughout the algorithm, this probability distribution is propagated unchanged for any SNP *x_i_* and receives no message 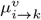 from its corresponding variable node.

##### Correlation Factor Nodes

We use LD to enhance the robustness of the system against correlation attacks. Hence, malicious service providers will not be able to use the SNPs-which are correlated with other SNPs with high probability-for watermark detection. For every SNP pair, correlation coefficients are calculated before the iteration and the pairs with coefficients higher than *σ_l_* threshold are marked as correlated and sensitive. Correlation coefficients may differ dependent on the states of each data point and their impact on estimating the probability distributions are typically asymmetric. For each sensitive SNP pair, there is one correlation node and these nodes keep track of the correlations inside the data.

The intuition for calculating the message to be sent by correlation node is derived from the definition of r-squared, or the coefficient of determination, that explains how good the proportion of variance in the dependent variable predicts the proportion of variance in the independent variable (Glantz *et al*., 2016). Since our system uses and infers marginal probability distributions in BP, we use 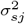 as a metric of how well we can predict the probability distribution of one state using the probability distributions of other correlated states (Miller, 1994). The messages from correlation node *c_s,j_* = *i* to *j^th^* variable node 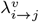 are calculated as

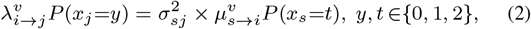

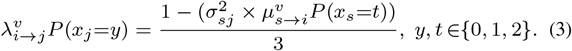

In these equations,

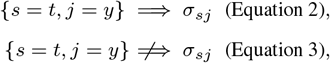

where *s* is the neighbor variable node, *σ_sj_* denotes the correlation coefficient, and 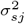 denotes the coefficient of determination. For further insight, please confirm the example in Appendix § 3.1.

##### Familial Factor Nodes

Familial factor node, *fam_i_* calculates message 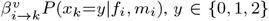 using the Mendelian Inheritance Law of Segregation and sends it to variable node *k*. Please confirm Appendix, Table 1 for Mendelian inheritance probabilities using the Law of Segregation. In the message, *f_i_* and *m_i_* corresponds to the *i^th^ SNP* values of father and mother, respectively.

For example, if the father has 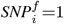 and the mother has 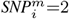 for the *i^th^ SNP*, the message from familial node *fam_i_* is as follows:

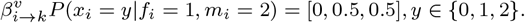

##### Phenotype Factor Nodes

Phenotype factor node, *phe_i_*, calculates message 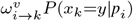, *y* ∈ {0, 1, 2}, *p_i_* ∈ {*dominant, recessive*} using the Mendelian Inheritance Law of Dominance and sends it to variable node *k*. Please confirm Appendix, Table 2 for Mendelian inheritance probabilities using the Law of Dominance. In the message, *p_i_* corresponds to the dominance trait of the observed phenotype in *i^th^ SNP*. *p_i_* can be either dominant or recessive. For example, if the data owner is known to have blue eyes (recessive gene), which is encoded in the *i^th^ SNP*, the message from phenotype node *phe_i_* is given as

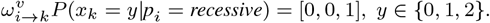

##### Attack-eLDP Nodes

The attack and *ϵ*-local differential privacy (*attack-eLDP*) node is designed to simulate the inference power of the attackers on data and calculates the inverse probabilities that will keep the attacker uncertainty at maximum against single SP and collusion attacks while keeping the local differential privacy criteria intact by updating the watermarked state options which violate *ϵ*-local differential privacy. This node receives a message from the variable node. Although acting as another factor node, it does not send the message to the variable node. Instead, the *attack-eLDP* node sends its message along with a variable node message to the watermarking algorithm as parameters.

Inside the *attack-eLDP* node, the attack part re-calculates the watermarking probabilities of all indices based on the variable node probability distributions and previously shared versions of states to simulate single SP and collusion attacks. In every *SP_k_*’s watermarking, a set of previous sharings 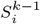 for each index *i* or set of indices I are used as a prior condition. Then the probabilities of the potential next states are calculated using binomial distribution given *S*. Finally, updated probability distributions are sent to the watermarking algorithm as the watermarking probability of each state. The calculation procedure followed by the node’s attack part is described in the sequel:

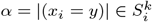, *α* is the number of states equal to *y* in set 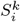.

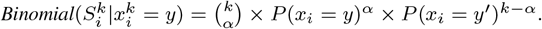
*P*(*x_i_*=*y*) and *P*(*x_i_*=*y*′) are calculated from the variable node’s message.

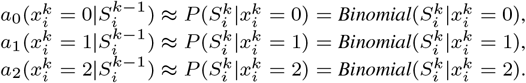

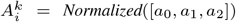, where *A* is the updated marginal watermarking probability distribution of *i^th^* index for the *SP_k_*.

The eLDP part, checks whether there are any states that violate the Local Differential Privacy (LDP) (Kairouz *et al*., 2014) condition for Alice who wants to have a plausible deniability factor for the versions of data she shares. Condition is satisfied if no state violates the Equation 3 in Appendix § 1.1. Probability distributions of the violating states converge by continuous averaging with variable nodes until non-violation. This incorporation creates watermarks for all the *SNPs* of the data owner and acts as a lower bound of privacy ensured along with lower and upper bounds on confidence degree by its very definition. For further insight, please confirm the example in Appendix § 3.2.

### 4.2 Watermarking Algorithm

*SNP* state inferences are assumed to be conducted by malicious SPs as well, given their prior information on the data for SP and Correlation Attacks (cf. § 3.3). We consider attacker inference strength and privacy criteria at the same time during watermarking where modifying the actual state of the data is mandatory. Modifying more indices than necessary results in losing utility on data. These modifications increase the detection probabilities of changed indices by malicious SPs and decrease efficiency. These modifications must be interpreted as actual data, not to give further means to malicious SPs for detecting watermarked indices. For example, watermarking a *SNP_i_* with *MAF_i_* = 0 is meaningless. Because no state of the *SNP_i_* other than homozygous major is observed, any change will be artificial and interpreted as watermarked.

Our watermarking scheme uses a probabilistic watermarking pattern rather than deterministic. To achieve this, we use a different set of indices and states to be watermarked for each SP. If we use fixed watermarked indices, it presents a risk of compromising watermark robustness against modifications and removals in single SP attacks and collusion attacks. If we use fixed watermarked states for each index, the data do not reflect the population distribution and using the probabilistic inference, attackers can identify the indices that show discrepancies with the population.

Given these criteria, we calculate a watermark score that helps us to list indices better to watermark in descending order. This score is calculated by comparing the *attack-eLDP* marginal probability distributions with the original states of data. Firstly, the probability of the actual state in *attack-eLDP* distribution is subtracted from one. This gives us the probability of that index being watermarked. Then these indices are sorted in descending order to give priority on indices most likely to be watermarked. For further insight, confirm the watermarking algorithm given in Appendix § 3.3.

## 5 Evaluation

We evaluated the proposed watermarking scheme in various aspects like watermark security against detection (robustness), the length of the watermark (utility loss) and the privacy guarantees. These aspects and their correspondence to the dependent variables are also given. We give the details of the data model, the experimental setup, and the results of the experiments in the sequel.

### 5.1 Data Model and Experimental Setup

For the evaluation, we used the SNP data of 1000 Genomes Project (IGSR, 2013). The data set contains the 7690 SNP-long data of 99 individuals in the form of 0s, 1s, and 2s, which is represented as a 99 × 7690 matrix. This data set is used for learning the linkage-disequilibrium and MAF statistics along with parental data generation based on the method proposed in (Deznabi *et al*., 2018). These statistics are then employed in the BP algorithm for probabilistic state inference. The threshold of pairwise correlations used for the results is specified as *ρ* = 0.9. Throughout the experiments, the length of data *d_l_* is fixed to 1000, the number of service providers (*h*) is fixed to 20, and *w_l_* values vary between 10 and 100. In exceptional cases, watermarks with *w_l_* > 100 are also tested, too.

### 5.2 Evaluation Metrics

We evaluated the data by calculating precision values and *ϵ*-local differential privacy achieved for various attack types, parameter configurations, and sets of predicted SPs. In collusion attacks, two SPs collusion scenario contains all 190 pairs of 20 SPs since, *h* is fixed to 20 and *C*(20, 2) = 190. This number increases rapidly as the number of collusion SPs increases. To keep the computational cost low, we took the number of malicious SPs scenarios as 190 unique random sets for each case. Besides, we kept the number of malicious SPs up to *k* = 10, since we assumed to know the *k* and *k* > 10 increased our detection results back. We find the malicious set of SPs by checking the watermark patterns. For the details of detection algorithms, please confirm § 4.1 in Appendix.

### 5.3 Results of Attacks

We evaluated the proposed scheme for the attack model described in § 3.3. The robustness of the watermarks is evaluated against single SP and collusion attacks in which the knowledge of single SP and correlation attacks are incorporated to reflect the worst-case scenario. In these experiments, we assume worst-case scenarios to create lower bounds. The assumptions that give maximum malicious SP information are as follows.

- Malicious SPs know the exact value of watermark length (*w_l_*).
- For every SP, *I_k_* is identical *k* ∈ {1, 2, …, *h*}. It means all SPs have the same set of indices of data.
- Malicious SPs have all the population information e.g. correlations, MAFs, frequency of states.
- Malicious SPs know the SNPs of the data owner’s father and mother.
- Malicious SPs know the phenotypical features of the data owner and correspondent SNP states.

#### 5.3.1 Single SP Attack

In single SP attacks, a single SP uses all the knowledge available to itself for inferring the marginal state probabilities of SNPs. This process is similar to the calculations in the belief propagation part of our watermarking scheme. Later on, malicious SP identifies the top *w_l_* SNPs with least probabilities *P*(*x_i_* = *y*), *y* ∈ {0, 1, 2} as watermarked, and modifies them to their most likely states for the prior information available to itself. One thing to note here is that these SNPs can be removed or partially modified as a variant attack scenarios. However, the total modification almost always yielded the best results in our experiments in favor of malicious SP and decreased our detection precision the most. Therefore, we assume the worst-case scenario against modifying attacks.

Figure 2 shows the impact of watermark length on precision for various values of *LDP* coefficients (*ϵ*). Impact of detection methods’ and *ϵ*s on single SP attack precision can be seen in Appendix § 4.2. In all cases, *w_l_* ≥ 40 seems to be the breaking point where the precision reaches to almost 100%. In our data, we think that this corresponds to the minimum amount of change needed for distinguishing a version of shared data from another effectively. For example, this *w_l_* may not suffice for comparison among 50 SPs. We observe that *ϵ* has no significant impact on precision.

**Fig. 2.**
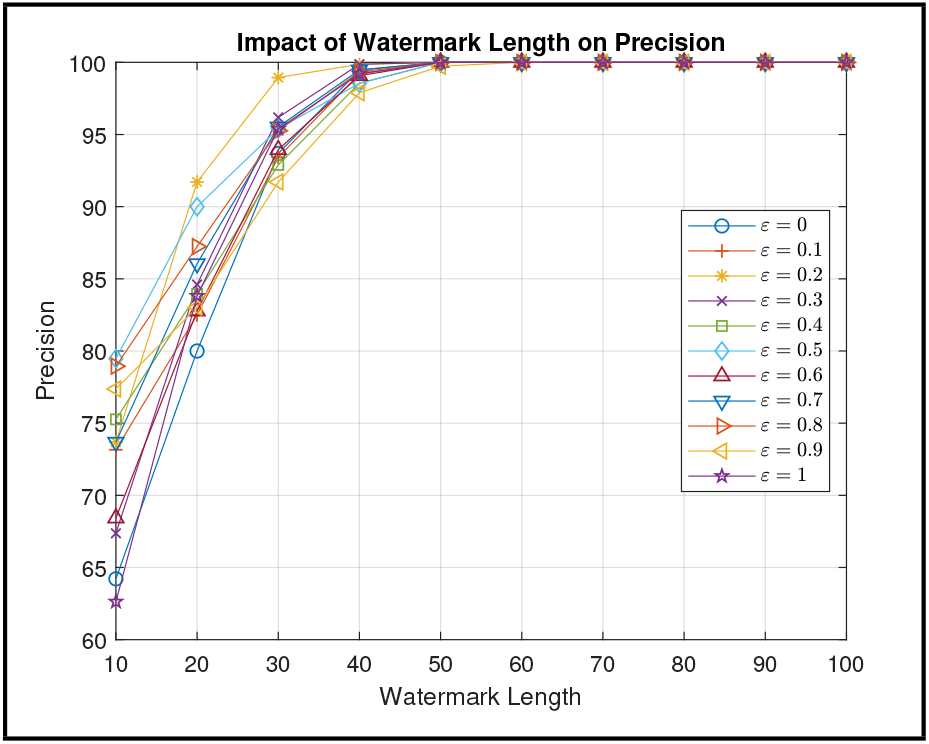
The impact of watermark length and *ϵ* on precision for a single SP attack.

#### 5.3.2 Collusion Attack

In collusion attacks, multiple SPs collude and bring their data together to detect and modify the watermarked indices. Firstly, the states different for the same SNP are identified as watermarked because the watermark pattern is unique for each SP. The maximum number of indices that can be identified as watermarked by malicious SPs is *w_l_* × *k* where *k* is the number of malicious SPs. Secondly, when the malicious SPs find fewer points in collusion attack than *w_l_* × *k*, they target additional indices as watermarked using the prior information on the data, e.g., *MAFs* and *LD* correlations, similar to the single SP attack. These indices are usually the least likely states when prior information is considered. At the end of the collusion attacks, malicious SPs change the states of data in two ways. The states that are not the same across all malicious SPs are modified to the most frequent ones. Then, the states that are the same across all malicious SPs but having the least likelihoods are modified to the most likely states possible. Our initial detection method, modification setups, and assumptions on the collusion attack are similar to those of a single SP attack. Since collusion attacks contain much more information about the probability of a state than the single SP attack, we expect the precision results of collusion attacks to be lower than single SP attacks. We expect a decrease in precision with the increasing number of malicious SPs.

Figure 3 shows the impact of watermark length on precision for *ϵ*=0. The impact of detection methods and *ϵ* on precision can be seen in Appendix § 4.3. Similar to single SP attacks, *ϵ* has no significant impact on precision. Among 20 SPs, *k* = 10 gives the worst results. Precision decreases as the number of malicious SPs increases but after *w_l_* ≥ 50 almost all *k* malicious SP scenarios are detected with precision rates higher than 80%. With *w_l_* = 100, precision increases up to 90% even for the worst case of 10 malicious SPs. We conducted experiments for *w_l_* > 100 and in some cases achieved precision up to 98% for *k* = 10.

**Fig. 3.**
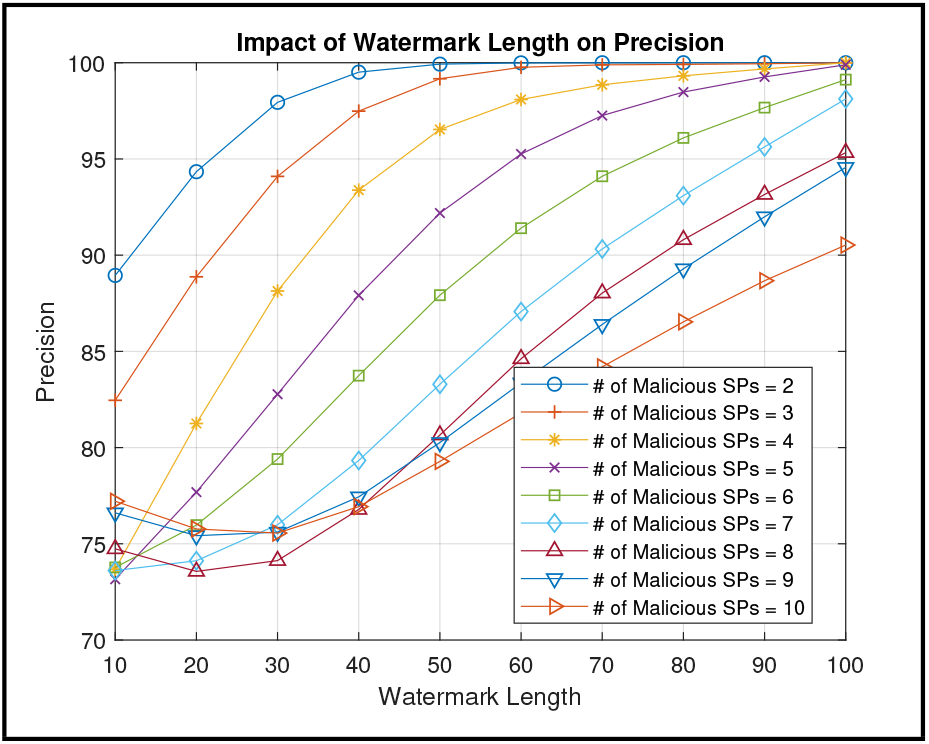
The impact of watermark length on precision for a collusion attack (*ϵ*=0).

## 6 Conclusion and Future Work

We propose a novel watermarking scheme for sequential genome data employing belief propagation (BP) with ensured *ϵ*-local differential privacy. This system is designed to be used between the data owner and service providers some of whom are assumed to be malicious. We implemented the algorithm against the worst-case scenario of malicious SPs. We assume that SPs know almost all statistics of the data. Therefore, we tested the robustness of watermarks against single SP attacks and collusion attacks. Unauthorized sharing risk is greatly mitigated by the BP algorithm. The algorithm secure high precision rates even against the worst-case scenarios when all potential prior information that malicious SPs can use are considered. We keep the changes on data minimum for preserving the utility of data by using short watermark length (*w_l_*). Our experiments show that when *w_l_* is kept higher than 50, even for a high number of malicious SPs, robustness is preserved more than 80%. We observe that *ϵ* does not significantly affect the precision. Privacy is preserved without impacting the precision and it addresses a potential liability issue from data being known and offers a privacy measure of plausible deniability needed especially for rare SNPs.

As non-ideal situations, the detection methods of the proposed algorithm do not predict but know the exact number of malicious SPs in the attack model. It is due to the very close scoring results of detection algorithms that reveals no clear-cut pattern and malicious SP identifications. Another such situation is that we consider the utility as the amount of data changed. Depending on the use-case, some SNPs may be considered to have more or less utility. As future work, the proposed scheme can be tested against larger datasets with more SPs. The impact of less powerful or inverse correlations on precision can be investigated. For addressing the non-ideal situations, an automated detection method with SP classifier that either diversifies or clusters malicious SPs from/with others by itself or a system that makes predictions like (Ayday *et al*., 2019) can be implemented. Finally, a utility quantification method can be implemented inside Belief Propagation as a factor node to assign priority on SNPs based on utility and alter the watermarking probabilities as intended.

