## Appendix for "Privacy-Preserving and Robust Watermarking on Sequential Genome Data using Belief Propagation and Local Differential Privacy"

### 1 Related Background

#### 1.1 Genomics

##### 1.1.1 Minor Allele Frequencies

The frequency of minor alleles prompts an immense amount of selection in heritability, and therefore, recorded as publicly available data for medical institutions (Hernandez *et al.*, 2019). Using these publicly available Minor Allele Frequency (MAF) values, the probability of each genetic state in populations can be inferred using the following equations:

$$AA = 0 : P(\textit{Homozygous Major}) = (1 - \textit{MAF})^2$$
$$Aa = 1 : P(\textit{Heterozygous}) = 2 \times (\textit{MAF}) \times (1 - \textit{MAF})$$
$$aa = 2 : P(\textit{Homozygous Minor}) = (\textit{MAF})^2$$

##### 1.1.2 Linkage Disequilibrium

*Linkage Disequilibrium (LD)* is the non-random heritable associations of alleles in different loci (Slatkin, 2008). Because it is an indication of population genetic forces on the genome formation, it is a widely investigated and exploited research topic in evolution and demographics studies (Stephens, 2001). Factors that have an impact on LD may vary due to genetic reshuffling, mutation rate, allelic drift, and so on. In genomic privacy, LD can be used to infer the state probabilities, and hence, the values of multiple Single Nucleotide Polymorphisms (SNPs) in correlated loci given the state value of a single SNP. Therefore, highly correlated states can be used for the enhancement of beliefs in belief propagation setup on the other SNPs. By exploring all the coexisting pairs of SNPs in the large sample population, linkage disequilibrium loci of high correlation values can be identified.

##### 1.1.3 Mendelian Inheritance Probabilities

Tables 1 and 2 present the Mendelian inheritance probabilities using the Law of Segregation and the Law of Dominance, respectively.

Table 1. Mendelian inheritance probabilities using the Law of Segregation.

| *SNP* Father (Rows) Mother (Columns) | 0 | 1 | 2 |
|---|---|---|---|
| 0 | [1, 0, 0] | [0.5, 0.5, 0] | [0, 1, 0] |
| 1 | [0.5, 0.5, 0] | [0.25, 0.5, 0.25] | [0, 0.5, 0.5] |
| 2 | [0, 1, 0] | [0, 0.5, 0.5] | [0, 0, 1] |

Table 2. Mendelian inheritance probabilities using Law of Dominance.

| Observed phenotype trait | *SNP* distribution |
|---|---|
| Dominant (AA or Aa) | [0.5, 0.5, 0] |
| Recessive (aa) | [0, 0, 1] |

#### 1.2 Local Differential Privacy

Differential Privacy is a system of public data sharing that uses the patterns of groups in the dataset and while doing so without compromising the privacy of individuals in the dataset (Dwork, 2006). The main intuition is that an algorithm is differentially private if the use of any particular individual's data cannot be inferred from the computations. If any inference probability is limited to the upper bound of $\rho < \epsilon$ in the dataset, the algorithm is $\epsilon$-differentially private. $\epsilon$-differential privacy is derived for a process $A$ if Equation 1 is satisfied in any two neighboring databases $D_1$ and $D_2$ with an outcome $O$.

$$P[A(D_1) = O] \leq e^{\epsilon} \times P[A(D_2) = O]. \quad (1)$$

Equation 1 is symmetrical and valid for any two neighboring databases $D_1$ and $D_2$. So, this equation can also be written as:

$$e^{-\epsilon} \times P[A(D_2) = O] \leq P[A(D_1) = O] \leq e^{\epsilon} \times P[A(D_2) = O]. \quad (2)$$

Local Differential Privacy (LDP) is the localized version of differential privacy that targets not the datasets or databases but the data indices. In LDP, the data is intentionally perplexed by the data owners so that plausible deniability is ensured without a "trusted party." The privacy assured by data owners is expressed as $\epsilon$-local differential privacy. This $\epsilon$ value can be thought to provide $\frac{100}{e^{\epsilon}+1}\%$ plausible deniability. As $\epsilon$ gets smaller, although the outcomes become less likely to be different from one another, a high level of privacy is ensured. In our use case, LDP is the local implementation and satisfying Equation 2 on every single data point or sequential data. We use LDP as an additional privacy preservation measure. It is a technology adopted by major technology firms like Google, Apple, and Microsoft for collecting mass anonymized data like web browsing behaviors, typing behaviors, and telemetry data (Cormode *et al.*, 2018).

A watermarking algorithm with $\epsilon$-local differentially privacy must normally satisfy Equation 2, for all sharings of all *SNPs*. This condition is used for limiting the amount of information gained by the exclusion of each shared data from the total set of sharings.

In Equation 2, the uncertainty increases with increasing $\epsilon$ at the expense of privacy. As $\epsilon$ decreases, more privacy is ensured. However, Equation 2 does not cover a localized setup but counts on databases or datasets as a whole. Therefore, we use a variant of differential privacy adapted from the geo-indistinguishability study of Andrés *et al.* (2012) that is both localized in *SNP* level and well suited to our sequential data. In e-DP part, our framework updates the probabilities of watermarking options that violate the local differential privacy condition of Andres et al. and his modified privacy criteria is given as

$$e^{-\epsilon\times r}\times\frac{P(x)}{P(x')}\leq\frac{P(x|S)}{P(x'|S)}\leq e^{\epsilon\times r}\times\frac{P(x)}{P(x')},\qquad \forall r>0,\quad \forall x,x':d(x,x')\leq r, \tag{3}$$

where $S$ represents the set of previous sharings of data and $r$ represents the distance between the states. In location data, $r$ is calculated as the maximum Euclidean distance between states. Since our data is sequential genomic data and the states have no priority over one another, we used Hamming distance for $r$, which is equal to one all the time. It is important to note that the first part of Equation 3 refers to the newly updated probabilities obtained from modifications such as adding and removing noise and it corresponds to the ratio between $ae_i$s. The second part refers to the unchanging probabilities of states given the prior information and it corresponds to the ratio between $var_i$s. We can compare the results for each $x_i = y, y \in \{0, 1, 2\}$, and decide whether the states should be updated or not for preventing violations against privacy condition.

### 2 Belief Propagation Algorithm

*Belief Propagation (BP)*, also known as sum-product message passing, is a message-passing algorithm used for the inference of networks and graphs like Bayesian Networks and Markov Networks (Braunstein *et al.*, 2005). BP calculates marginal probability distributions of unknown variables in factor graphs in an iterative manner using the information from previous states. In a factor graph, two types of nodes are used: (i) Factor Nodes, and (ii) Variable Nodes. BP is a widely used technique in graphs because the marginal probability computation of variables that have a dependency on multivariate data (factors) gets exponentially complex as the number of factors increases. Moreover, marginal probabilities of factors must be re-computed given the new distribution of variables. With a finite number of iterations, BP approximates the actual distribution with a low computational cost.

The steps of the proposed BP-based algorithm are as follows:

- The algorithm starts in a variable node with an initial probability distribution.
- The algorithm collects messages from the factor nodes for updating the probability distributions of the targeted unknown variable nodes. In loopy bilateral graphs, this process is handled in iterations until convergence. However, this approach is changed to a top-to-bottom approach with one or two iterations for tree-like graphs like ours for efficient approximation.
- Variable nodes generate the factor node messages by multiplying all incoming messages from the neighbors except the receiver neighbor.
- Factor nodes generate the messages by using local functions and send them to corresponding variable nodes.
- At the end of each iteration, the marginal probability distribution of each variable node is updated by multiplying all incoming messages from neighbors.
- The algorithm approximately calculates the beliefs of the variable nodes and passes it to the "Attack and $\epsilon$-local differential privacy" (*Attack-eLDP*) node.
- The *Attack-eLDP* node acts as a secondary factor node and calculates a new message that considers both attack scenarios and local differential privacy criteria.
- Finally, the *Attack-eLDP* node passes its message together with variable node messages as parameters into the watermarking algorithm.

### 3 The Details About the Nodes Used in Belief Propagation Algorithm

#### 3.1 Example From Correlation Node

Figure 1 shows how correlation nodes are connected with variable nodes and how they send messages to one another.

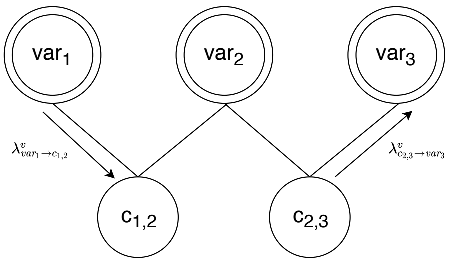


**Fig. 1.** The relationship between variable nodes and correlation nodes. Both nodes may receive and send messages. For simplicity, one message for each type is shown.

For example, $SNP_1$ is connected with $SNP_2$ via $c_{1,2} = i$ for $x_1 = 0$, $x_2 = 2$ with $\sigma_{1,2} = 0.9$ and $\mu^v_{1\to i}P(x_1 = y) = [0.3, 0.6, 0.1]$, $y \in \{0, 1, 2\}$ at the $v^{th}$ iteration. Then, we may calculate the message from $i$ to $2^{nd}$ variable node, $\lambda^v_{i\to 2}P(x_2 = y)$ as:

$$\lambda^v_{i\to 2}P(x_2=y)=\left[\frac{1-(0.9^2\times 0.3)}{3},\ \frac{1-(0.9^2\times 0.3)}{3},\ \frac{1-(0.9^2\times 0.3)}{3}\right]+\left[0,0,(0.9^2)\times(0.3)\right],\left.\right]$$

$$\lambda^v_{i\to 2}P(x_2=y)=[0.2523, 0.2523, 0.4954].$$

#### 3.2 Example From Attack-eLDP Node

For example, assume for the index $i$, $var_i = \mu^v_{i\to ae_i}P(x_i{=}y) = [0.6, 0.4, 0]$ sends the following message to the *attack-eLDP* node $ae_i$ and $S_i^{k-1} = \{0, 0, 1, 0, 1, 0\}$ where $k = 7$. We may calculate the watermarking probabilities of index $i$ for $SP_7$ as follows:

For $x_i^7 = 0, \alpha = 5 \Longrightarrow$
$\textit{Binomial}(S_i^7|x_i^7 = 0) = \binom{7}{5} \times (0.6)^5 \times (0.4)^{7-5} = 0.261,$
For $x_i^7 = 1, \alpha = 3 \Longrightarrow$
$\textit{Binomial}(S_i^7|x_i^7 = 1) = \binom{7}{3} \times (0.4)^3 \times (0.6)^{7-3} = 0.290,$
For $x_i^7 = 2, \alpha = 1 \Longrightarrow$
$\textit{Binomial}(S_i^7|x_i^7 = 2) = \binom{7}{1} \times (0)^1 \times (1)^{7-1} = 0.$

$A_i^7 = \textit{Normalized}([0.261, 0.290, 0]) \approx [0.474, 0.526, 0]$ is the updated watermarking probability distribution of index $i$ for $SP_7$.

Continuing from the above example, we can calculate the privacy conditions as follows:

$$\text{For } x_i^7=0,\ \frac{P(x|S)}{P(x'|S)}=\frac{0.474}{0.526}=0.901,\ \text{and}\ \frac{P(x)}{P(x')}=\frac{0.6}{0.4}=1.500.$$

This means $e^{-\epsilon} \times 1.500 \leq 0.901 \leq e^{\epsilon} \times 1.500$ must be satisfied for not violating the $\epsilon$-local differential privacy. Since $\frac{0.901}{1.500} \leq 1$ and $ln(0.601) = -0.509$, if $\epsilon \leq 0.509$ the state probability of $P(x_i^7 = 0)$ must be updated for not violating the privacy by averaging the attack node probabilities with variable node probabilities. After $P(x)$ is set, the distribution is continuously normalized to converge into the probability that satisfies the condition.

$$\text{For } x_i^7 = 1, \ \frac{P(x|S)}{P(x'|S)} = \frac{0.526}{0.474} = 1.110, \text{ and } \frac{P(x)}{P(x')} = \frac{0.4}{0.6} = 0.667.$$

This means $e^{-\epsilon} \times 0.667 \leq 1.110 \leq e^{\epsilon} \times 0.667$ must be satisfied for not violating the $\epsilon$-local differential privacy. Since $\frac{1.110}{0.667} = 1.664$ and $ln(1.664) = 0.223$, if $\epsilon \leq 0.223$ the state probability of $P(x_i^7 = 1)$ must be updated likewise.

$$\text{For } x_i^7 = 2, \frac{P(x|S)}{P(x'|S)} = \frac{0}{1} = 0, \text{ and } \frac{P(x)}{P(x')} = \frac{0}{1} = 0.$$

This means $e^{-\epsilon} \times 0 \leq 0 \leq e^{\epsilon} \times 0$ and it never violates the local differential privacy for any $\epsilon$ just like the case of $x_i^7 = 0$. However, we know that $P(x_i = 2) = 0$ and it is an impossible watermarking case regardless of the violation.

If Alice set her privacy criteria to $\epsilon < 0.509$, $ae_i$'s final marginal probability distribution will be equal to the distribution enforced by the eDP part. Otherwise, the distribution remains the same as attack part determined, which is equal to $[0.474, 0.526, 0]$.

#### 3.3 Watermarking Algorithm

**Algorithm 1** Watermarking Algorithm

```
Watermark(data, atks, vars, wScore, w_l)
j ← 1
k ← 0
newdata ← data
while k < w_l do
  i ← wScore(j, 4) {index of SNP}
  temp ← data(i) {actual state of SNP}
  flag ← true
  while flag do
    r ← random(0, 1)
    if atks(i, 1) ≥ r then
      newdata(i) ← 0
      if temp ≠ 0 then
        k + +
      end if
      flag ← false
    else if atks(i, 1) + atks(i, 2) ≥ r then
      newdata(i) ← 1
      if temp ≠ 0 then
        k + +
      end if
      flag ← false
    else if atks(i, 1) + atks(i, 2) < r then
      newdata(i) ← 2
      if temp ≠ 0 then
        k + +
      end if
      flag ← false
    end if
  end while
  if j == length then
    j ← 1
  else
    j + +
  end if
end while
return newdata
```

### 4 Evaluation

#### 4.1 Detection Methods

For detection, we compare the attacked data produced by malicious SPs with each of our previously shared data and watermark patterns. Assuming, we know the number of malicious SPs, we use two detection methods *Hamming Distance (H)* and custom *spPenalizer (E)* and their relaxed versions *Hamming Distance Relaxed (HR)* and *spPenalizer Relaxed (ER)*. $H$ method as the name suggests, calculates the Hamming distances between attacked data and every shared SP data by checking every index. After the distances are calculated, all SPs are sorted from least different than the attack data (least distance) to the most different (most distance) and top $k$ (# of malicious SPs) SPs are flagged as malicious. $E$ method, firstly identifies the indices that has not been the same with the original data at least once (conflicting indices) for all SPs. Then, the variance of each conflicting index is calculated as a penalizing factor. Depending on the match of indices between shared SP data and attack data, penalizing factors are added into the matching SPs. Finally, top $k$ SPs who are penalized the most are flagged as malicious SPs. In the relaxed versions of both methods, top $k$+1 are used in flagging process.

### 4.2 Single SP Attack

Figure 2 shows a comparison of the detection methods for a single SP attack. We observe that the relaxed versions *HR* and *ER* outperform the methods *H* and *E*. However, the methods *E* and *ER* outperform the methods *H* and *HR*.

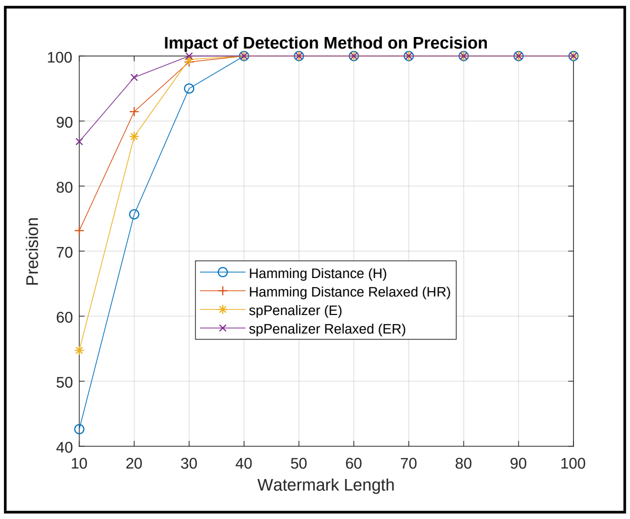


**Fig. 2.** The impact of watermark length and detection method on precision for a single SP attack ($\epsilon$=0).

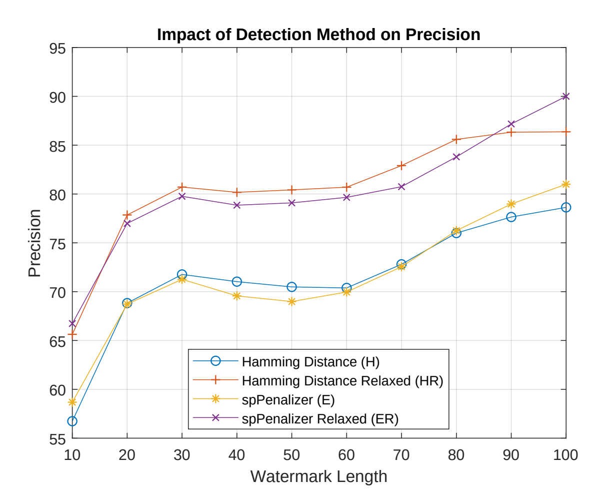


**Fig. 3.** The impact of watermark length and detection method on precision for a collusion attack ($\epsilon$=1).

### 4.3 Collusion Attack

Figure 3 shows a comparison of the detection methods for a collusion attack. Similar to single SP attack, we observe that the relaxed versions *HR* and *ER* outperforms the *H* and *E* methods. Differing from the single SP attack though, the results of *H* and *E* and *HR* and *ER* are almost identical. Figure 4 shows that the relaxation of the privacy criteria $\epsilon$ to 0.5 or 1 does not impact the precision results at all.

Figure 5 depicts the impact of privacy criteria ($\epsilon$) on precision. The case for $k = 2$ provides very limited information because all of the $w_l$, except $w_l = 10$ gives 100% precision. For both $k = 6$ and $k = 10$ though, the precision remains stable with naturally lower precision results for $k = 10$. Sometimes, arbitrary decreases and increases occur throughout different watermark lengths but overall, $\epsilon$'s little to no impact can be observed here as well.

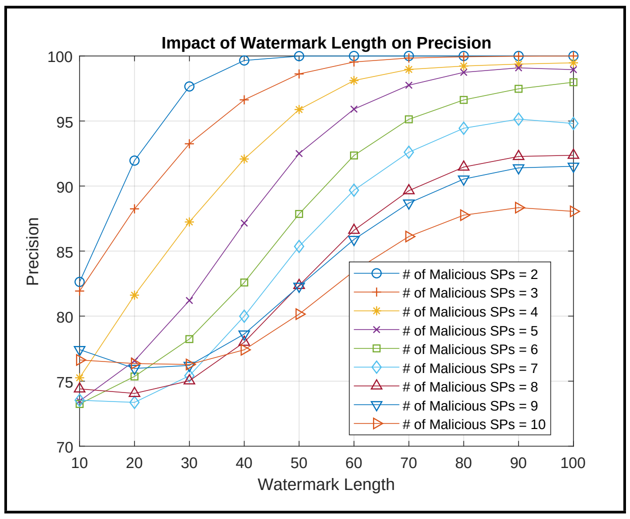


(a) $\epsilon = 0.5$

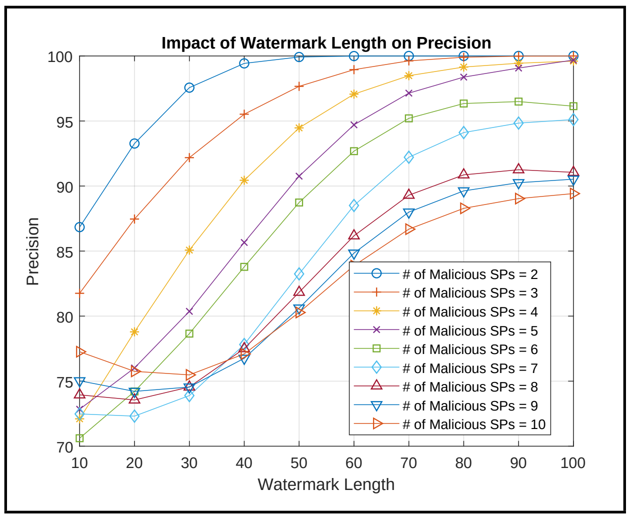


(b) $\epsilon = 1$

**Fig. 4.** The impact of watermark length on precision for collusion attacks with different number of malicious SPs and $\epsilon$s.

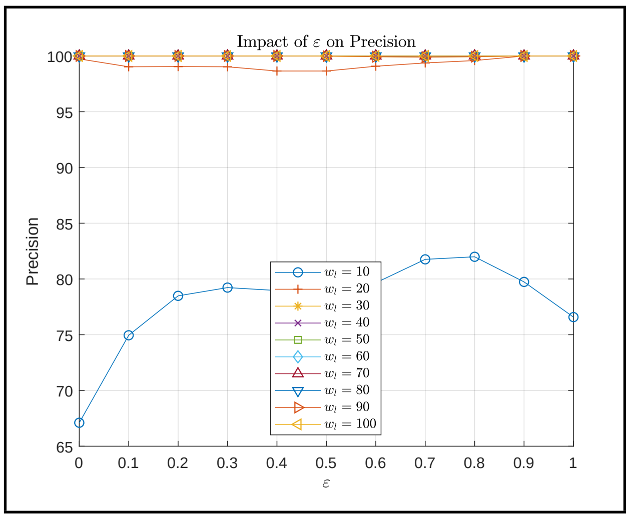


(a) $k = 2$

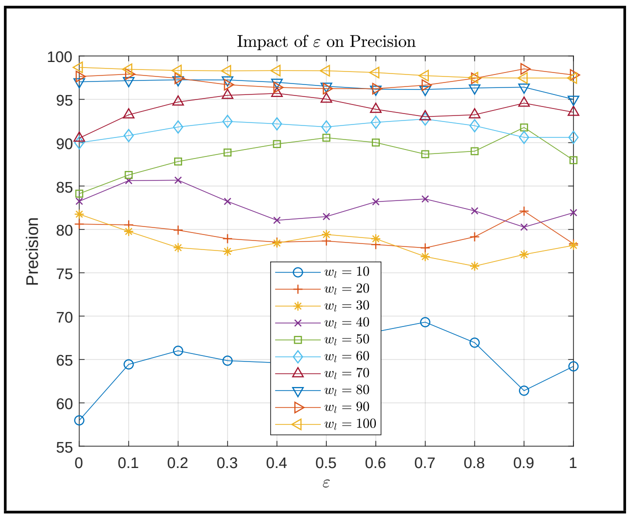


(b) $k = 6$

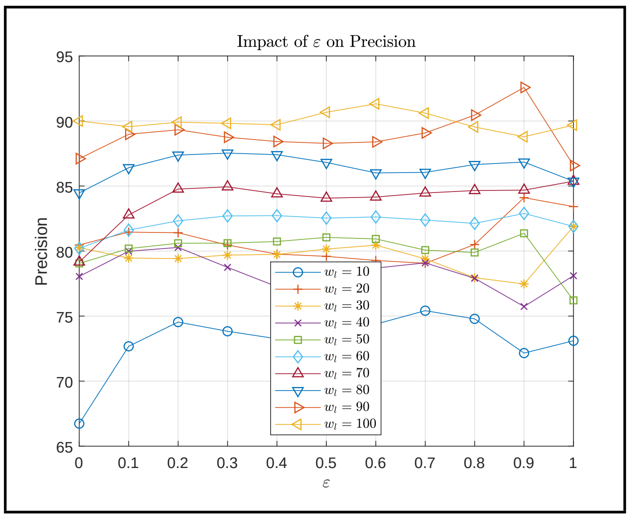


(c) $k = 10$

**Fig. 5.** The impact of $\epsilon$ on precision for collusion attacks with different number of malicious SPs and $\epsilon$s.
